# Defining the role of *Pseudomonas aeruginosa* PilY1 in signaling and virulence

**DOI:** 10.1101/2025.05.15.654335

**Authors:** Christopher L. Pritchett, F. H. Damron, M. Barbier

**Affiliations:** College of Public Health, East Tennessee State University, Johnson City, TN 37614; Department of Microbiology, Immunology, and Cell Biology, West Virginia University, Morgantown, WV 26506-9600

## Abstract

*Pseudomonas aeruginosa* (Pa) is an important opportunistic pathogen that has many virulence factors expressed in a coordinated manner to cause infection. External appendages, such as the type IV pili (T4P), can be used for signal transduction changing gene expression using a mechanism termed mechanosensing. An important role for the minor pilin, PilY1, is to signal that the bacterium has attached to a surface such as a host cell. This mechanosensing is important for triggering changes in bacterial gene expression allowing colonization of host tissues. In this study, we identify new PilY1-controlled genes and demonstrate the importance of PilY1 in controlling second messengers, including cAMP. Genetic analysis showed that PilY1 is important in controlling cAMP levels by regulating the expression of the main adenylate cyclase, *cyaB*. PilY1 also controls the expression of PA4781 a putative cyclic di-GMP phosphodiesterase. PilY1 also controls the expression of the *algZ/R* operon by preventing autoregulation of this system. Furthermore, the minor pilin PilY1 functions in different *P. aeruginosa* strains, including mucoid strains, demonstrating conservation of function. Surprisingly, bacterial survival in the lung, liver, and blood did not require PilY1. Overall, these findings suggest that PilY1 is critical for regulating second messengers by preventing the AlgZ/R system from functioning. Our work provides insight into the mechanisms that *P. aeruginosa* uses to modulate second messengers which are important in biofilm formation and virulence.

Importance. PilY1, a component of type IV pili, is important for signaling environmental cues into the bacterial cell. However, the signaling pathway PilY1 controls is poorly understood. We determined that PilY1 is involved in the regulation of cAMP. Transcriptomic analysis identified new PilY1-controlled genes. We also demonstrate the functionality of PilY1 in other strains, including mucoid strains commonly found in chronic infections. We elucidate the importance of PilY1 controlling cAMP by repressing the AlgZ/R system and preventing *cyaB* expression. Our data help to explain how regulatory systems interact in *P. aeruginosa* providing insight that may be useful for therapeutic development.

## Introduction

*Pseudomonas aeruginosa* is a human opportunistic pathogen capable of causing fatal infections in immunocompromised individuals such as those undergoing chemotherapy, those suffering from severe burn wounds, chronic obstructive pulmonary disease (COPD), or cystic fibrosis (CF) (1–3). Intrinsic and acquired drug resistance has made *P. aeruginosa* a microbe to fear in the clinical setting (4, 5). In the disease process, the adherence to host tissues is a prerequisite for colonization and the T4P play a dominant role in adhesion (6–9). Upon inhalation or aspiration, *P. aeruginosa* uses the T4P to colonize the respiratory tract by allowing movement along epithelial cells and forming microcolonies to initiate infection (9). There is some evidence that the T4P bring the bacterial cell in close contact with host cells allowing the type III secretion system to inject effectors and penetrate deeper into host tissues (7, 10). Recent work has discovered that the T4P are also important sensors for determining when the bacteria have contacted a surface such as a host cell (11, 12). Understanding the molecular events that transpire during these initial events can inform the development of new treatment methodologies to combat *P. aeruginosa* infection.

The T4P is made up of the major pilin subunit PilA, and the minor pilins FimU-PilVWXEPiY1lY2 (13–16). PilY1 is the tip adhesin of the T4P and functions in twitching motility, adherence, and mechanosensing (7, 17–20). PilY1 is part of the operon with the other minor pilins and this operon is controlled by the transcriptional regulators Vfr and the AlgZ/R two-component system (21–24). The AlgZ/R system is comprised of the membrane histidine kinase AlgZ and the response regulator AlgR which activate transcription of numerous genes (25). Both Vfr and AlgR are important for virulence (26, 27) but their contribution to virulence in terms of PilY1regulation is not known.

Once the pilin components are assembled into mature T4P, the retraction and extension of pili could be used for motility across cellular surfaces or allow tight binding to host cells and this is critical for virulence (9). PilY1 is also important for regulating pilin secretion and is therefore also important for T4P extension and retraction (28). The T4P are also important for biofilm formation (29). The T4P are involved in many different aspects of *P. aeruginosa* biology, making this appendage an attractive drug target.

PilY1 signaling impacts second messengers with one mechanism for regulating c-di-GMP known. PilY1 senses surfaces upregulating c-di-GMP levels to increase biofilm formation (11, 19, 30). In contrast, PilY1 was shown to be important for upregulating cAMP upon surface binding (12, 19). However, no mechanism for how PilY1 controls cAMP is known. Other studies have demonstrated that cAMP and c-di-GMP are inversely correlated (31, 32) so how PilY1 affects both c-di-GMP and cAMP has not been clarified. If PilY1 were to increase cAMP levels, then this could explain decreased virulence of the *pilY1* mutants in various pathogenesis models (33, 34) due to decreased virulence gene expression due to Vfr inactivity. Vfr is activated by cAMP leading to increased production of several virulence factors including the type III secretion system (12, 23). Additionally, cAMP-Vfr inhibits *fleQ* expression (35). FleQ is a regulator of *cdrA* expression commonly used to monitor c-di-GMP levels (36). FleQ bound to c-di-GMP activates *cdrA* expression enhancing biofilm formation in *P. aeruginosa* (37). The lack of a specific mechanism for how PilY1 regulates secondary messengers, such as cAMP, requires further study.

The full extent of genes PilY1 controls is not known. Determining the other players affected by PilY1 is important to understanding the signaling network PilY1 controls and could provide new targets to combat *P. aeruginosa*. We hypothesized that PilY1 controls factors involved in cAMP metabolism. Using RNAseq, we discovered that PilY1 controls cAMP levels through at least two different mechanisms. First, PilY1 decreases cAMP production by preventing AlgZ/R activation of the main adenylate cyclase, *cyaB*. Second, PilY1 controls the expression of PA4781 that acts as a second cAMP phosphodiesterase. Altogether, our results indicate that PilY1 function is critical for proper secondary messenger levels and suggest a model for how PilY1 influences the activity of transcriptional regulators in *P. aeruginosa*.

## Materials and Methods

### Bacterial strains, plasmids and growth conditions

Bacterial strains and plasmids used in this study are listed in Table 1. *P. aeruginosa* was grown at 37°C in Luria Bertani (LB) broth supplemented with 50mM MOPS or LB agar for routine growth. Transcriptional fusion assays were performed in LB Mops (50mM), quorum sensing media (QSM, 22mM KH_2_PO_4_, 22mM Na_2_HPO_4_, 85mM NaCl, 10% tryptone, 1 mM MgSO_4_, 0.1 mM CaCl_2_). Media used for *P. aeruginosa* was supplemented with tetracycline (180 μg/ml), gentamicin (250 μg/ml), carbenicillin (300 μg/ml), or trimethoprim (500 μg/ml) as needed. *E. coli* was cultivated at 37°C in LB and supplemented when necessary with ampicillin (100 μg/ml), kanamycin (35 μg/ml) or chloramphenicol 34 μg/ml. Yeast-tryptone media (YT, 1% tryptone, 0.5% yeast extract) was used for allelic exchange experiments and was supplemented with the appropriate antibiotics or with 10% sucrose. In the case of complementation, spectinomycin (25 μg/ml) was used in combination with trimethoprim (500 μg/ml).

### Mutant Strain construction

PCR generated fragments were amplified using Q5 polymerase (New England Biolabs) and cloned directionally into either pEX18Tc or pEX18Gm (38) after performing cross-over PCR (39). Mutant constructs were introduced into PAO1 or other strains via tri-parental mating using the helper strain pRK2013 (40). Single crossover mutants were selected on the appropriate selective media and merodiploids were grown without selection to select for a second crossover event. Mutants were plated on YT 10% sucrose and then patch-plated on selective media and PIA. Colonies not growing on selective media were screened via PCR for the appropriate mutation. Complementation was accomplished by PCR of the corresponding wild-type gene using Q5 (NEB) and cloning into the integrating vector pTJ1 (41). Sequencing was used to confirm all constructs and mutant strains.

### RNA seq

Total RNA was isolated from the selected strains using the RNA SNAP procedure (42) and DNase-treated. The RNA was purified using RNeasy kit (Qiagen), quantified using Nanodrop ND-1000 (Nanodrop), and assessed for RNA integrity using Agilent BioAnalyzer RNA Pico chip (Agilent). A total of 50 μg from each sample was sent for RNAseq analysis (Admera Health). All samples were submitted to rRNA depletion and reassessed for RNA integrity. rRNA depleted mRNA samples were then fragmented and prepared into libraries using Illumina TruSeq RNA library prep kit v2 (Illumina). The libraries were then sequenced on an Illumina HiSeq 2 × 150 bp reads with a total of 10^6^ reads per sample and 3 replicates per experimental condition. Sequencing data were deposited to the GEO website and are available under the reference number GSE278651. RNA-seq reads were analyzed using the software CLC Genomics workbench (Qiagen). The *P. aeruginosa* PAO1 genome and annotations were downloaded from the Pseudomonas Genome Database and reads were mapped using the following settings: mismatch cost = 2, insertion cost = 3, deletion cost = 3, length fraction = 0.8, similarity fraction = 0.8. RPKM values were generated using default parameters for CLC Genomics. Fold changes in gene expression and statistical analyses were performed using an Extraction of Differential Gene Expression (EDGE) test with the Bonferroni correction.

### Transcriptional fusion analysis

Upstream DNA fragments containing promoter regions were generated by using primers listed in Table 1 in conjunction with Q5 polymerase (New England Biolabs, Ipswich, MA). PAO1 genomic DNA was used as template. PCR products were cloned into pMiniT (NEB) and then sub-cloned into miniCTXlacZ or miniCTXlux using the restriction enzymes *Hind*III*/Bam*HI, *Hind*III/EcoRI, or *Kpn*I/*Bam*HI (NEB). PCR products were purified, cut with restriction enzymes and inserted into the EcoRI/BamHI sites of miniCTXlacZ using T4 DNA ligase (NEB). Strains were selected for tetracycline resistance and then conjugated with pFLP2 to remove vector sequences in the case of *lacZ* fusions (38). Strains were selected for carbenicillin resistance, grown overnight without selection and plated on YT media with 10% sucrose to select for the loss of pFLP2. Individual colonies were patch-plated onto VBMM CB300 and PIA to ensure the loss of pFLP2. To confirm the presence of the fusion constructs, PCR was performed using the forward primer used to construct the fusion and the reverse primer, lacZRforTF (Table 1). β-galactosidase activity was determined by incubating cell extracts with ONPG (4mg/ml) as per Miller (43). A strain carrying the empty vector, miniCTXlacZ, was also conjugated into PAO1, assayed and this background (28 Miller Units) was subtracted from all transcriptional fusions. All mucoid strains were confirmed mucoid at the end of each experiment by plating on PIA plates to ensure all colonies were mucoid. All assays were performed in buffered LB (LB + 50mM MOPS). At least three biological replicates were reproduced for all assays.

For *lux* fusions, promoter fragments were cloned into miniCTXlux and conjugated into *P. aeruginosa* strains (44). The vector sequences were not removed. White-walled plates in a Synergy HTX (Bio Tek) were used for assays. Strains were grown overnight, diluted 1:50 in LB MOPS (50mM) and 200 μl was used to inoculate white-walled plates containing LB MOPS (50mM). Assays were measured for 5-10 hours depending on the strains used. Data was analyzed using GraphPad Prism. All experiments were repeated at least three times in triplicate for each strain. Data is reported as relative luminescence by dividing luminescence readings by the OD_600_ for luminescence assays and in Miller Units for β-galactosidase assays. Error bars indicate the standard error of the mean.

### AlgR Purification and antibody production

The *algR* gene was cloned into the expression vector pMALc6t expression vector (NEB). AlgR was expressed as a fusion with maltose-binding protein (MBP). The fusion protein was batch purified using agarose resin and cleaved using TEV protease and the TEV and MBP removed using Ni-affinity chromatography. AlgR was dialyzed using a slidalyzer (ThermoFisher) and Storage Buffer (20% glycerol, 20mM Tris pH=7.5, 5mM MgCl_2_, and 1mM DTT) overnight at 22°C. The purity of AlgR was visually determined in a Coomassie stained 4-12% gradient electrophoresis gel (SDS-PAGE) and AlgR was confirmed by western blots using AlgR-specific sera. Antibodies were produced to AlgR by ProSci (Poway, CA).

### Electrophoretic mobility shift assays

Gel mobility shift assays as described previously with some modifications (45). PCR fragments were generated using biotinylated primers (see Table 1) and gel purified using the Monarch Gel Extraction kit (NEB). Binding reactions were carried out using purified AlgR. The DNA (50ng-200ng) probes were mixed with AlgR protein containing 20 mM Tris-HCL (pH 8.0), 0.5 mM dithiothreitol, 20 mM KCL, 0.5mM MgCl_2_, 2 mM EDTA and 5% glycerol. The nonspecific competitor poly (dI-dC) was added at 10 μg/ml for all gel shift reactions. After incubation for 20 minutes at room temperature (25°C), the samples were separated by electrophoresis on a 6% native polyacrylamide gel with 0.375 X TBE used as running buffer for approximately 1.5 h at 100 Volts. Purified AlgR was incubated at increasing concentrations to determine suitable concentrations to be used. The gel shifts were developed using the chemiluminescent detection kit (Life Technologies) and visualized using a Bio-Rad chemi-doc. Gel shifts were repeated at least three times, and a representative gel shift is shown.

### Site-directed mutagenesis

Primers are listed in Table S1. The primers were phosphorylated and used in site-directed mutagenesis as per the manufacturer’s instruction using Q5 (NEB). Constructs were analyzed by restriction enzyme analysis and sequencing. Mutant strains were constructed using homologous recombination as described above and were checked using PCR and primer pairs in Table S1. Site-directed mutants had an *Eco*RI site engineered to allow easier detection of the promoter mutation. Additional PCR amplicons were sequenced to confirm the mutation in each strain using the same primers. Further confirmation of mutants was done using phenotypic or biochemical assays.

### Western blot analysis

The bacteria were collected by centrifugation and resuspended in 50 mM Tris-HCl pH 7.5 and lysed using sonication or B-PER (Fisher Scientific). Total protein concentrations were quantified by the Bradford protein assay (Bio-Rad). Cell extracts (5-10 μg) were separated by SDS-PAGE on 10% polyacrylamide gels and transferred to a polyvinylidene difluoride membrane (GE Osmonics). The membranes were probed using a 1:10,000 dilution of anti-AlgR rabbit polyclonal antibody followed by a 1:20,000 dilution of horseradish peroxidase-conjugated goat anti-rabbit monoclonal antibody (46) and the signal was detected using chemiluminescence (Bio-Rad). The anti-AlgR polyclonal antibody was pre-absorbed with cell extract from the *ΔalgR* strain to remove other non-specific antibodies.

Westerns were developed using ECL reagent (ThermoScientific) and imaged using a Chemi-Doc (Bio-Rad).

### Phenotypic Assays

Pyocyanin assays were performed as described in (47) . Briefly, overnight cultures in QSM media were pelleted and the supernatant filtered through a .2μm filter. Three ml’s of supernatant was extracted twice with 1ml of chloroform and vortexed vigorously. The organic layer was removed and 1 ml of 0.2N HCl was added. The OD_520_ was taken and the samples normalized to the OD_600_ of the culture. Blanking of the spectrophotometer was done using 0.2 N HCl. For congo red binding, congo red plates were inoculated with 2 μl of overnight cultures after washing with sterile PBS. Swarming assays were performed as described previously (48). Biofilm assays were performed as described previously (49).

### Competition experiments

All animal protocols were approved by the Institutional Animal Care and Use Committee at East Tennessee State University (P220301) following the guidelines of the Office of Laboratory Animal Welfare. Overnight cultures were diluted 1:100 in LB and grown for 3-5 hours, pelleted, and washed 2 times with PBS. The OD_600_ of each strain was adjusted to an OD_600_= 0.2. Further dilution with PBS was done to achieve the final dose of 1 x 10^7^ CFU’s.

Mutant and wild-type strains were mixed 1:1 and the input determined by plating on PIA and PIA gent. Groups of five 12-16 week CD-1 mice (Envigo) were infected by oropharyngeal aspiration after anesthetization with 2.5% isofluorane with *P. aeruginosa* at doses of 1 x 10^7^ CFU’s in 40 μl. Sixteen hours after infection, mice were euthanized and blood, lungs, livers, and spleens were collected. Organs were homogenized in 1 ml of PBS and all tissues were serially diluted and 100 μl plated in duplicate on both PIA and PIA Gent. Competitive indices were determined by dividing the output ratio by the input ratio.

## RESULTS

### PilY1 controls cAMP levels

PilY1 is important for *P. aeruginosa* virulence (7, 11, 12, 33). However, a mechanism for attenuation has not been uncovered. One consequence of mechanosensing is the increase in cyclic di-GMP (20). Notably, cAMP levels inhibit c-di-GMP levels in *P. aeruginosa* (31, 32, 35). This led us to speculate that PilY1 might impact cAMP levels in *P. aeruginosa* and this could explain decreased c-di-GMP in the *pilY1* mutant. Transcriptional *lux* reporters were constructed to monitor cAMP and c-di-GMP production over time. We integrated the second messenger sensors into the *P. aeruginosa* chromosome to analyze the transcriptional fusions in single copy (50, 51). A *cdrA* transcriptional fusion using the *lux* reporter system was constructed and used as a sensor for c-di-GMP as this promoter has been shown to correlate with c-di-GMP levels (36). We detected a decrease in c-di-GMP levels in Δ*pilY1* (Fig. 1A). A Δ*gacA* strain was used as a negative control (52). The *cdrA* reporter in a Δ*pilA* strain had no difference from the wild-type PAO1 strain. A *ΔpilJ* strain was also similar in c-di-GMP levels to the wild-type strain. These data suggest that PilY1 has a unique function in controlling c-di-GMP levels compared to other pilin components tested.

**Figure 1.**
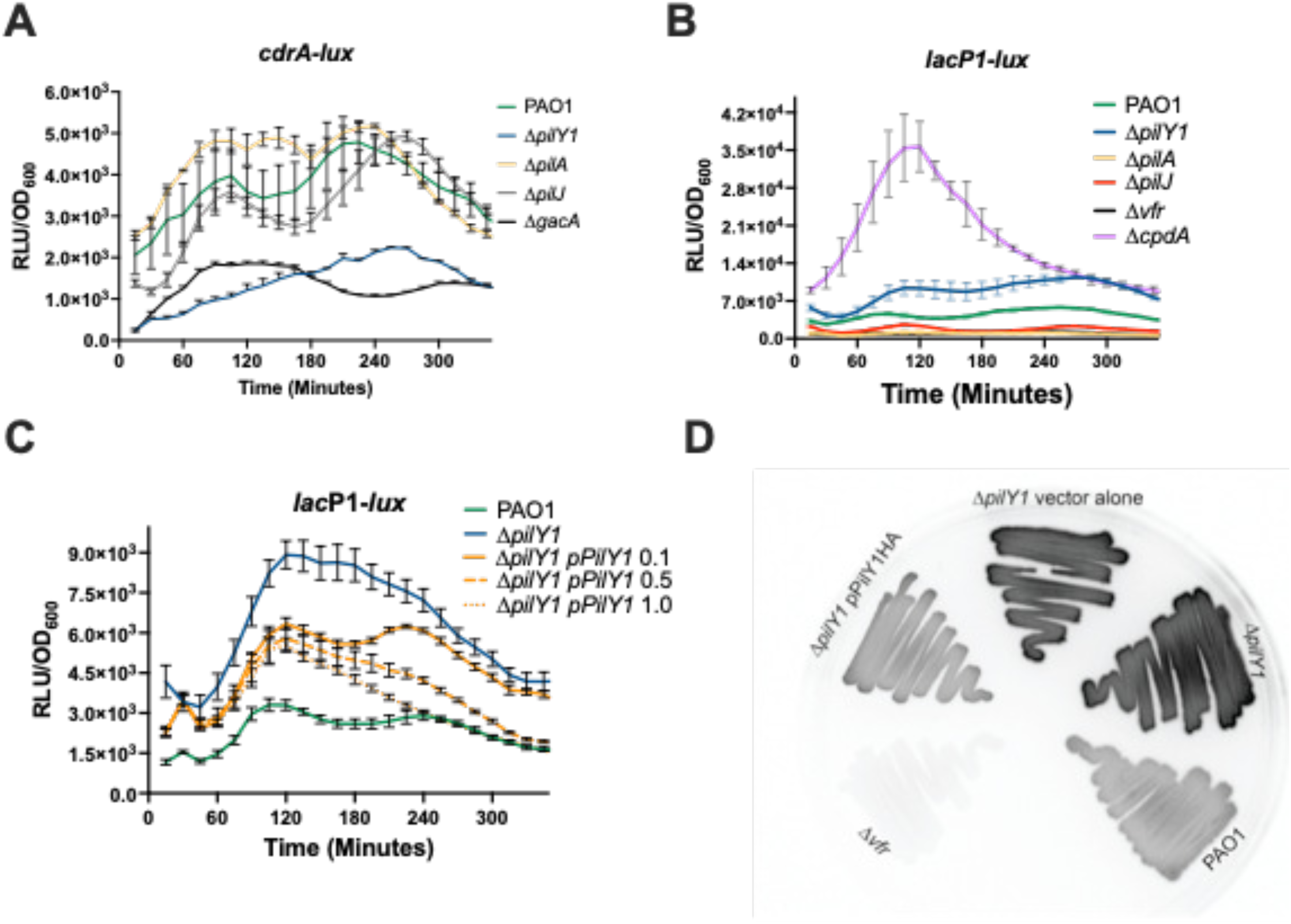
PilY1 regulates both cAMP and c-di-GMP in *P. aeruginosa*. A. *cdrA* promoter activity representative of c-di-GMP levels. B. *lac*P1-lux reporter activity representative of cAMP levels. C. Complementation of Δ*pilY1* reduces cAMP. D. Luminescence of strains containing the *lac*P1-lux reporter on solid media after 16 hours. Assays represent three biological replicates done in triplicate and measurements were taken every 15 minutes for five hours. Complementation studies were with L-arabinose (0-1%).

The *ΔpilY1* mutant was complemented by incorporating a C-terminal HA-tagged version of the wild-type *pilY1* using an integrating vector to provide single copy complementation (41). This construct was fully functional as determined by complementing twitching motility and biofilm formation (FigS1). When Δ*pilY1* was complemented, there was delay in the response, but increasing arabinose concentration restored *cdrA* reporter activity (Fig S2). In fact, increasing arabinose to 0.5 and 1.0% resulted in increased *cdrA* reporter activity after 180 minutes suggesting that increasing the amount of PilY1 leads to increased cyclic di-GMP (FigS2). This further supports the importance of PilY1 in controlling c-di-GMP levels.

A cAMP *lux* reporter using the *lacP*1 promoter previously shown to correlate with cAMP levels (19, 31, 53, 54) was used as a readout for cAMP levels in pili mutants (Fig 1B). A Δ*vfr* mutant was used as a negative control as this strain is not responsive to increased cAMP levels (53). We also tested the cAMP reporter in a Δ*cpdA* mutant as a positive control (55), which had the highest luminescence. A Δ*pilY1* mutant had modestly increased cAMP levels compared to PAO1 using the *lac*P1-lux reporter in a liquid assay (Fig 1B). We also tested the *lac*P1-*lux* reporter in other pili mutants to determine if the increased cAMP was specific to *pilY1* mutants or if this was a general phenomenon of pili mutants. Previous work indicated that both PilA and PilJ were necessary for optimal cAMP levels after surface attachment (56). A Δ*pilA* strain had decreased Lux activity compared to PAO1 and was like the Δ*vfr* strain. A Δ*pilJ* strain had slightly decreased luminescence compared to PAO1. Therefore, our results are consistent with published results for Δ*pilA* and Δ*pilJ* (12, 19, 53) and indicates our reporter is functioning as expected. Complementation of the *ΔpilY1* strain containing the *lac*P1-*lux* reporter demonstrated decreased cAMP levels even without induction with arabinose (Fig 1C). Increasing arabinose concentration lowered cAMP reporter activity indicating that PilY1 is responsible for lowering cAMP levels. Complementation using 0.5 or 1% arabinose reduced cAMP to wild-type levels in the complemented strain after 240 minutes. From these data, we conclude that PilY1 regulates cAMP levels in *P. aeruginosa*.

Because the levels of luminescence detected in the liquid assay were small, we decided to test the reporter strains under different conditions. A previous study found that cAMP levels are increased upon growth on surfaces (19). The strains containing the *lac*P1-*lux* reporter were grown on solid media and produced luminescence that was visually striking. The luminescence of the *ΔpilY1* was much brighter than the wild-type and the *Δvfr* mutant (Figure 1D).

Complementation with *pilY1* restored the level of luminescence to the wild-type strain (Fig 1D). These data suggested that PilY1 is important for decreasing cAMP levels in *P. aeruginosa* and that PilY1 differs from other pili components in terms of controlling cAMP.

### Transcriptomic analysis reveals unknown PilY1-controlled genes

RNAseq was performed to provide an unbiased view of what other genes PilY1 might control and to determine how PilY1 might be involved in pathogenesis. Comparing the Δ*pilY1* mutant to the wild-type strain PAO1, RNAseq identified 168 genes that were upregulated in the *ΔpilY1* strain and 118 that were downregulated (Figure 2, Supplementary Tables S1 and S2). PilY1 is known to control pyocyanin (57) and control expression of its own operon (34). As expected, many downregulated genes were involved in pyocyanin production (Fig. 2). Twelve of the downregulated genes (Table S1) were from both operons involved in pyocyanin biosynthesis (58). The finding of pyocyanin biosynthetic genes validates our use of RNAseq to identify new PilY1 targets.

**Figure 2.**
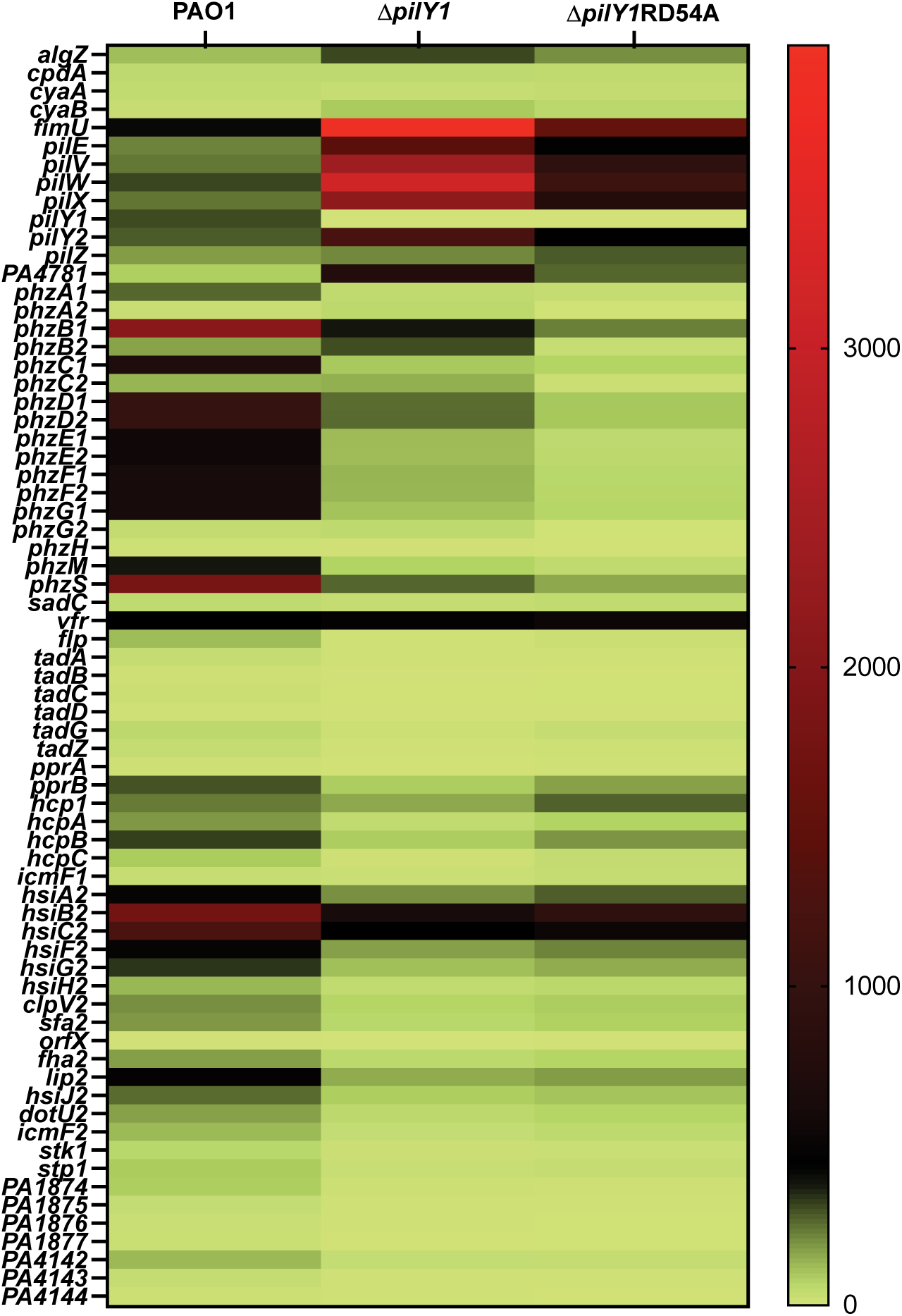
RNAseq analysis defines the PilY1-controlled genes. A. Heat map comparing differentially regulated genes. The heat map shows the reads per kilobase per million reads (RPKM) for 150 differentially expressed genes between the wild-type PAO1 and the Δ*pilY1* strains.

Other downregulated genes included several secretion systems. Two Type I secretion systems (PA1874-PA1877 and PA4142-4144) that are poorly understood were decreased in the *pilY1* mutant. PA1874-1877 is an ABC transport system that has been shown to be an efflux pump when *P. aeruginosa* is growing as a biofilm (59). Even less is known about the putative secretion cluster PA4142-4144, but it is regulated via quorum sensing (60). However, both type I secretion systems have been increased in expression in burn wounds (61). In addition, the second type VI secretion system (H2-T6SS (62)) was also decreased in expression when *pilY1* was deleted.

Interestingly, PilY1 appears important in the expression of type IVb pili. *P. aeruginosa* is one of very few bacteria that expresses both type IVa and type IVb pili (63). Several mRNA’s encoded by the *tad* locus and *flp* were severely decreased as well as the *pprAB* system that controls the type IVb structural genes. The type IVb pili are important for DNA transfer (64). Therefore, the RNAseq analysis suggests that there is a hierarchy in the expression of the type IVa and type IVb pili in *P. aeruginosa* with the type IVa pili controlling type IVb expression.

Upregulated genes include *fimU*, *pilX*, *pilW*, *pilE*, and *pilY2* contained within the *fimU* operon that also contains *pilY1* (Fig. 2, Table S2). Numerous unannotated genes were also increased in expression. Interestingly, the major adenylate cyclase, *cyaB* (23), was increased in the *pilY1* mutant. Overall, our RNAseq results identified new PilY1-controlled targets and suggested a possible mechanism for PilY1 control of cAMP levels.

Another goal was to determine other possible explanations for PilY1 control of c-di-GMP or cAMP. There was no decreased expression of any known diguanylate cyclase, but we did find one putative phosphodiesterase that had increased expression: PA4781 (Fig 2, Tables S1 and S2). An increase in PA4781 could explain the decreased c-di-GMP levels in addition to PilY1 affecting SadC (20, 65).

*P. aeruginosa* controls cAMP levels by two adenylate cyclases, CyaA and CyaB, and one known phosphodiesterase *cpdA* (23, 55). The RNAseq data did not indicate an increase in the main cAMP phosphodiesterase *cpdA.* We confirmed this result using a transcriptional reporter to the *cpdA* promoter and found that *cpdA* expression as slightly reduced (Fig S3A). However, we did find an increase in the major adenylate cyclase *cyaB* in the RNAseq analysis. Therefore, the RNAseq results suggested that increased cAMP was due to increased *cyaB* expression and not increased expression of the phosphodiesterase that breaks down cAMP. Overall, our RNAseq data identified new genes regulated by PilY1. Additionally, the data suggests that pyocyanin and other secretion systems are the main virulence factors regulated by PilY1.

### The major adenylate cyclase, *cyaB*, is increased in the Δ*pilY1* strain

CyaB provides most of the cAMP in the *P. aeruginosa* cell (23). We constructed a *cyaB*-*lux* transcriptional fusion to monitor *cyaB* expression kinetically and detected increased reporter activity in the Δ*pilY1* strain (Fig 3A). This result confirms the RNAseq data. We complemented the Δ*pilY1* strain containing the *cyaB*-*lux* fusion (Fig 3B). As little as 0.1% arabinose decreased expression of *cyaB*-*lux* in the complemented strain and all further arabinose increases had a similar effect (Fig 3B). From these data, we conclude that PilY1 does control the expression of *cyaB*.

**Figure 3.**
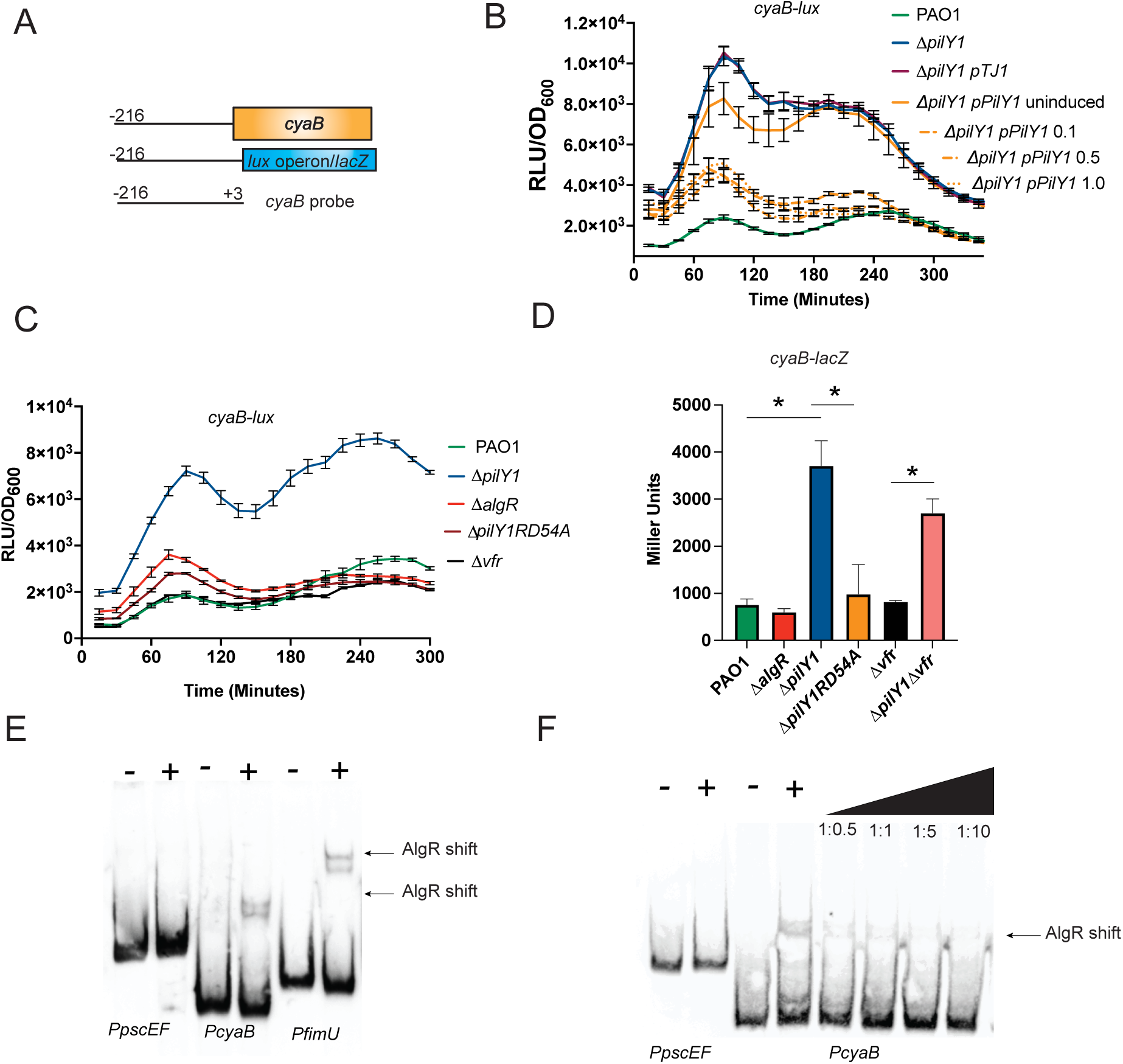
PilY1 controls *cyaB* using the AlgZ/R system. A. Schematic showing the region of the *cyaB* promoter used for constructing transcriptional fusions and making the probe for RMSA analysis. B. A *cyaB-lux* transcriptional fusion has increased activity in the Δ*pilY1* strain and overexpression of *pilY1* leads to decreased *cyaB* expression. C. The *cyaB-lux* in strains demonstrates the AlgZ/R system is important in *cyaB* expression. D. A *cyaB-lacZ* in strains confirms AlgZ/R dependence and the increased expression in the *pilY1* mutant. E. EMSA analysis of the *cyaB* , *pscEF*, and *fimU* probes using purified AlgR. F. Competition using unlabeled *cyaB* probe demonstrates the specificity of AlgR binding to *cyaB* promoter. Competitions were done using 0.5:1,1:1, 1:5, and 1:10 ratios of unlabeled to labeled probe, respectively. (*, *p*<0.001)

To determine if PilY1 affects the expression of other genes encoding proteins that make or degrade cAMP, transcriptional *lux* reporters were constructed of *cyaA* and *cpdA* (Fig. S3). Mutation of *pilY1* did not affect transcription of *cyaA* or *cpdA*. We also tested a Δ*vfr* mutant as Vfr and cAMP are intimately linked and Vfr has been demonstrated to directly activate *cpdA* expression (55). Compared to the wild-type strain, only *cyaB* differed significantly in expression of the adenylate cyclase genes in the *pilY1* mutant (Fig 3C and S3). Interestingly, Vfr did not affect the expression of *cyaB*, but *cyaA* was slightly increased in a *vfr* mutant at later timepoints (Fig S3B). There was a slight decrease in *cpdA* expression in the *pilY1* mutant: however, when complemented, the activity of the reporter was not restored to wild-type levels (data not shown). There was also decreased expression of *cpdA* in the Δ*vfr* strain used as a control (55) (Fig S3A). Altogether, these data suggest that Vfr only plays a significant role in regulating the phosphodiesterase *cpdA* and not the adenylate cyclases, *cyaA* and *cyaB*. Based on these data, PilY1 controls the expression of the major adenylate cyclase, *cyaB*. However, we cannot rule out that there is increased CyaB activity as well.

### AlgZ/R is responsible for increased cAMP levels in a *pilY1* mutant

A previous study demonstrated that two regulators, Vfr and the AlgZ/R two-component system, are increased in activity in a *pilY1* mutant (19). We investigated the role of Vfr and the AlgZ/R system in *cyaB* regulation. As shown in Figure 3C, there was decreased *cyaB* expression in a double mutant containing a deletion in *pilY1* and encoding a phosphorylation-incompetent version of AlgR compared to the Δ*pilY1* strain suggesting that phosphorylated AlgR is required for the increased *cyaB*. There was little change in the *cyaB* reporter expression in the Δ*algR* or the *Δvfr* strains compared to the wild-type strain. Even the wild-type strain had very little *cyaB*-*lux* reporter activity. These results suggest that phosphorylated AlgR, and not Vfr, regulate *cyaB* expression.

We constructed a second *cyaB* reporter using *lacZ* to allow quantitation of the increased expression and to confirm the *lux* reporter data. The Δ*pilY1* strain had a > 3-fold increase (P< 0.0005) in *lacZ* expression compared to the wild-type confirming the *lux* fusion data and the RNAseq data (Fig 3D). Once again, introducing the allele encoding a phosphorylation-incompetent version of AlgR into the Δ*pilY1* strain (Δ*pilY*1RD54A) abrogated the increased reporter activity (Fig 3D). A Δ*vfr* strain did not differ from PAO1, the wild-type strain. When *vfr* was deleted in the Δ*pilY1* mutant (Δ*pilY1*Δ*vfr*) there was almost a 3-fold increase in *lacZ* activity indicating that Vfr does not increase *cyaB* expression and consistent with the *cyaB*-*lux* data (P<0.0006) (Fig 3D). Additionally, the Δ*pilY1*Δ*vfr* strain did not differ significantly from the Δ*pilY1* strain (Fig 3D). These data indicate that AlgR, and not Vfr, is responsible for the increased *cyaB* expression in the Δ*pilY1* mutant.

To further support AlgZ/R regulation of *cyaB*, we constructed a Δ*algZ/R* strain that was complemented with the *algZ/R* operon under arabinose control. While no significant difference was found between PAO1 and Δ*algZ/R*, overexpression of the algZ/R operon resulted in increased *cyaB-lacZ* reporter activity (Fig S4). These data further support the role of AlgZ/R in control of *cyaB* expression.

A putative AlgR-binding site was found in the *cyaB* upstream sequence (Fig 4A). To determine if AlgR directly controlled *cyaB* expression by binding this sequence, purified AlgR (0.1μM) was tested with the *cyaB* promoter using gel shift analysis (Fig 3E-F). As a negative control, the *pscEF* genes were used. As a positive control the *fimU* promoter (which regulates *pilY1*) was also included. A shift comparable to the *fimU* positive control was evident (Fig 3E). Competition experiments using unlabeled *cyaB* as a probe confirmed that the binding of AlgR to the *cyaB* promoter was specific (Fig 3F). The gel shift studies indicated that AlgR directly binds the *cyaB* promoter.

**Figure 4.**
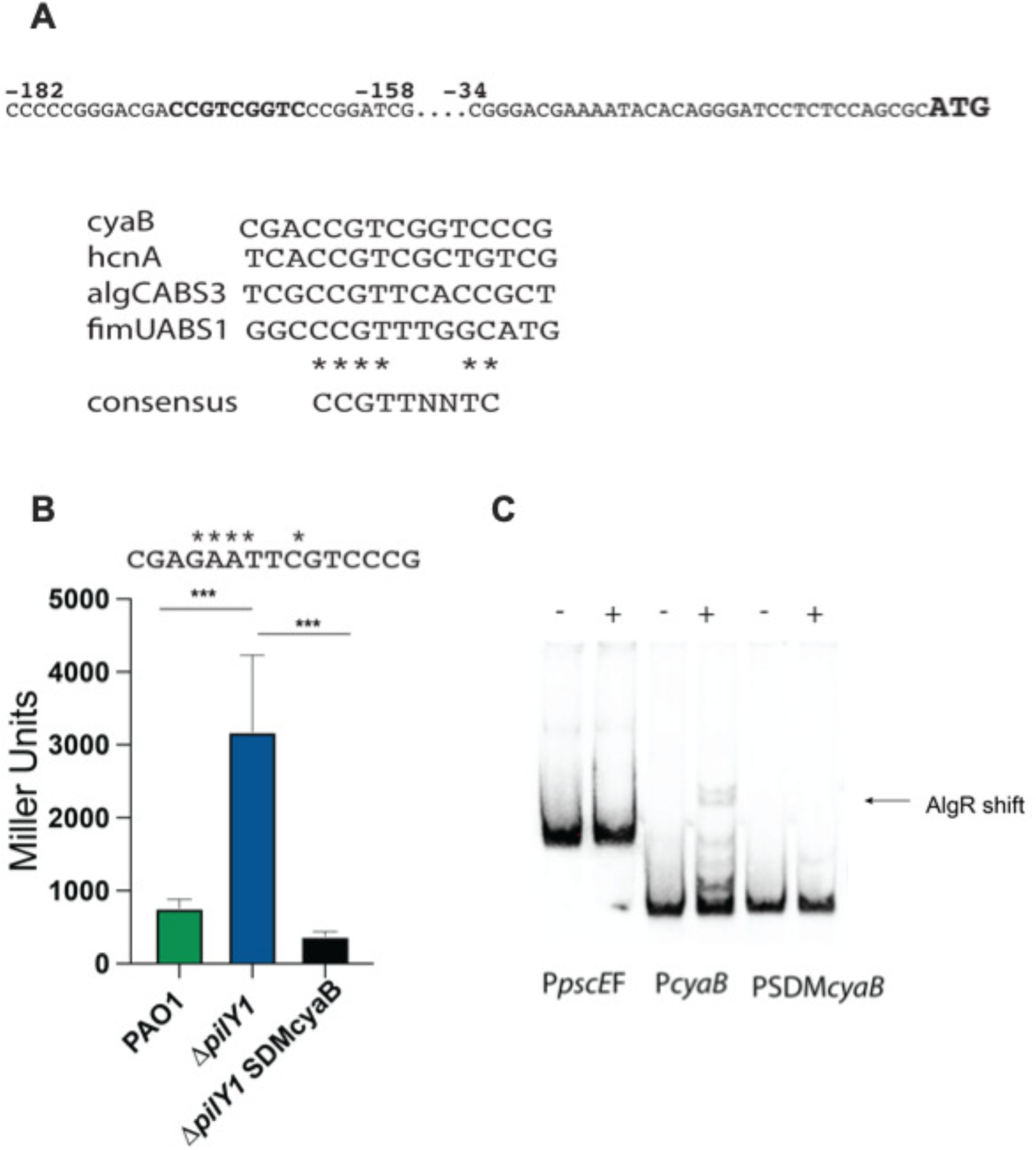
Site-directed mutagenesis of the AlgR-binding site of the *cyaB* promoter. A. Sequence of the *cyaB* promoter region. The putative AlgR-binding site is indicated in bold. Below is an alignment of the putative AlgR-binding site compared to other AlgR binding sites. The consensus sequence is shown below. B. Transcriptional fusion analysis of the *cyaB* promoter after site-directed mutagenesis. At the top indicates the sequence changes made to the putative AlgR-binding site. The strains and the allele of the *cyaB* promoter are located with the strains. C. EMSA demonstrating the importance of the single AlgR-binding site in the *cyaB* promoter. (***, *p*<0.001)

Inspection of the *cyaB* promoter identified a putative AlgR-binding site 170-162bp upstream (Fig 4A). The putative binding site only deviated at one nucleotide from the consensus sequence “CCGTTNNTC” (25) (Fig 4A). We used site-directed mutagenesis to mutate the AlgR-binding site into the *cyaB*-*lacZ* transcriptional fusion (Fig 4B). Once again, *cyaB* expression was more than 3-fold higher in activity in the Δ*pilY*1 mutant. However, the site-directed mutant had a greater than 4.5-fold reduction from the Δ*pilY*1 parent strain. Interestingly, mutation of the AlgR binding site also reduced the reporter activity to approximately half of the wild-type strain PAO1 but was not significant (Fig 4B).

Gel shift analysis was used to further investigate the role of the putative AlgR-binding site. We compared the native *cyaB* promoter to the site-directed mutagenized promoter. The native promoter bound AlgR as previously shown (Fig 4C). However, the mutagenized promoter was unable to bind to AlgR (Fig 4C). These data clearly demonstrate that AlgR directly regulates the *cyaB* promoter and identify the specific nucleotides involved. Therefore, the AlgZ/R system is important in the regulation of cAMP levels in *P. aeruginosa* by controlling *cyaB* expression. Overall, our data indicate that PilY1 prevents AlgZ/R activity leading to decreased *cyaB* expression decreasing cAMP levels.

To determine the biological relevance of AlgR regulation of *cyaB*, we constructed a mutant in the Δ*pilY1* mutant that contains an *Eco*RI site in place of the AlgR-binding site. This also aided detection of the promoter mutants during construction. We first tested the *lac*P1 and *cdrA* fusions to determine if AlgR control of *cyaB* might explain the dysregulation of second messenger production. In the case of *lac*P1-*lux*, the Δ*pilY*1SDM strain had the expected lower *lac*P1-*lux* reporter activity compared to the Δ*pilY1* parent strain (Fig S5A). In the case of the *cdrA*-*lux* reporter, the Δ*pilY1*SDM differed from their respective parent strain (Fig S5B). Overall, these results indicate that AlgR activation of the *cyaB* promoter is responsible for increased cAMP levels in the Δ*pilY1* strain, but do not explain the decreased c-di-GMP levels in Δ*pilY1* (Fig. S5B)

Previous work demonstrated that a Δ*pilY1* mutant had reduced pyocyanin (57). Confirming the previous study, we found decreased pyocyanin production in the Δ*pilY1* strain compared to PAO1 (Fig S5C). Pyocyanin was not significantly affected in the Δ*pilY1*SDM mutant compared to Δ*pilY1*. While AlgR control of the *cyaB* promoter is not responsible for the decreased pyocyanin in the Δ*pilY1* strain, this regulation is important in allowing proper pyocyanin production in the wild-type strain.

### PilY1 functions in multiple strain types, including in mucoid strains

Given the diversity of *P. aeruginosa* strains, we wanted to assess the functionality of PilY1 in other backgrounds. We used another strain PA103 (66) as this strain is similar to clinical strains causing high rates of pneumonia and sepsis (67–71). We also used clinical strains, strain 383, and its mucoid derivative 2192 (72), and PDO300, the *mucA* mutant of PAO1. A *fimU-lacZ* transcriptional fusion was used to assess the functionality of PilY1 in these strains as previous studies have shown that *pilY1* mutation led to increased expression of its own operon (19, 34). A *ΔalgR* strain was used as a negative control (22) as the AlgZ/R system positively regulates the *fimU* operon (22). As expected, there was a >20-fold increase in *fimU* expression in Δ*pilY1* versus PAO1 (Fig 5A). Analysis of strain PA103 (66), also resulted in increased *fimU* expression (P<0.0001) when *pilY1* was deleted in this background (Fig 5A). There was a 7-fold difference between the 383*ΔpilY1* strain and the parental strain 383 for *fimU* expression (P<0.0001) (Fig 5A). Overall, PilY1 functions in multiple *P. aeruginosa* backgrounds by repressing the AlgZ/R system.

**Figure 5.**
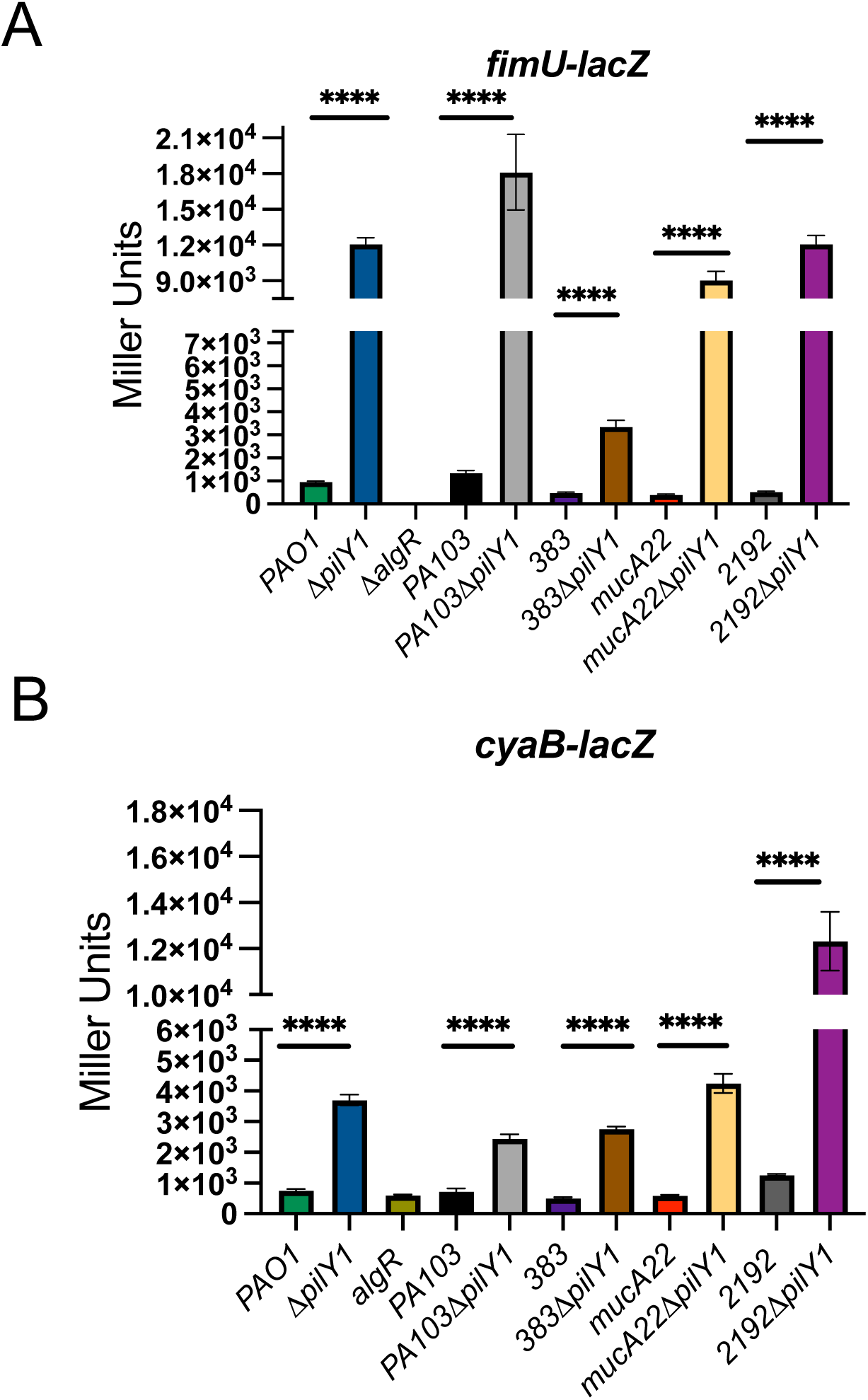
PilY1 functions similarly in both laboratory and clinical of *P. aeruginosa* strains. A. Beta-galactosidase activity of a *fimU* reporter. B. Betagalactosidase activity of a *cyaB* reporter in multiple *P. aeruginosa* strains. Transcriptional fusions were performed in triplicate at least three times. *p*<****, 0.001 based on Student9s t-test comparing the parent and the mutant strain.

In cystic fibrosis patients, *P. aeruginosa* frequently loses motility and persists as a biofilm (73). Whether pili components, such as PilY1, are produced in mucoid strains has not been investigated. The rationale for these experiments was that PilY1 in a chronic *P. aeruginosa* isolate had significant phenotypic changes (57), suggesting that PilY1 can function independently of being incorporated into the pilus. The laboratory strain PDO300 (derived from PAO1) and a clinical isolate 2192 (74), which both contain mutations in *mucA*, were used.

Deletion of *pilY1* in either *mucA* mutant background (PDO300 or 2192) did not significantly affect alginate production (data not shown). The *fimU* reporter had increased activity (> 9-fold) in the *pilY1* mutants in both PDO300 and 2192 (Fig 5A). These results suggested that PilY1 does function in *mucA* mutants.

While PilY1 controls *fimU* (and *pilY1*) operon expression (19, 34), the role of PilY1 and the AlgZ/R system controlling *cyaB* expression has not been explored. To determine if PilY1 was also decreasing *cyaB* expression in other backgrounds, the *cyaB*-*lacZ* transcriptional fusions was tested. There was increased activity of the fusion whenever a strain had been deleted for *pilY1* (Fig 5C). Both PAO1*ΔpilY1* and 383Δ*pilY1* had increased *cyaB* reporter activity compared to their respective parent strains (Fig 5C). We also saw an increase in *cyaB-lacZ* activity in the mucoid *ΔpilY1* strains in both genetic backgrounds (Fig 5C). When *pilY1* was deleted in PDO300 (PDOΔ*pilY1*), there was a ∼7-fold increase in *cyaB* reporter expression. In the case of 2192Δ*pilY1* there was an ∼ 10-fold increase (Fig 5C). This further confirms PilY1 is functional in *mucA* mutants and indicates that the signaling cascade is found in more than just the laboratory strain PAO1. These data also support that a major role of PilY1 is to repress the AlgZ/R system to prevent *cyaB* expression.

### PilY1 controls *algZ/R* expression

A previous study indicated that the AlgZ/R system was hyperactive in a Δ*pilY1* mutant and this was responsible for attenuation in a *C. elegans* model (34). Our RNAseq data indicated increased *algZ* expression in the Δ*pilY1* strain. Autoregulation is a common characteristic of two-component systems (75), but has not been described for the *algZ/R* system. The *algZ/R* operon is under complex genetic regulation where promoters lie upstream of *algZ* and within the *algZ* coding region (76)(Fig. 6A). At least one of the promoters upstream of *algZ* (Promoter 1) is regulated by Vfr (21, 76). The promoters within the *algZ* coding region controlling only *algR* expression are activated by the alternative sigma factor AlgU/T (76). To begin to investigate autoregulation of the *algZ/R* promoters, we constructed *algZ* and *algR* transcriptional fusions to separate the two promoter regions (Fig 6A) and assayed the activity in the minor pilin mutant Δ*pilY1*. Confirming the RNAseq data, the *pilY1* mutant had increased activity of the *algZ* transcriptional fusion, but there was no increased activity of the promoter located within the *algZ* coding region that controls only *algR* expression (Fig. 6B and C). If the minor pilin mutant has increased AlgZ/R activity, then this suggests that AlgR should exist in a phosphorylated state (77). When a double mutant consisting of the phosphodeficient allele of *algR* in the Δ*pilY1* background (Δ*pilY*1RD54A) was analyzed with the *algZ* transcriptional fusion, the reported luminescence returned to wild-type levels (Fig 6B). The transcriptional fusion analyses indicate that AlgR activates the expression of the *algZ*/*R* operon, but only from the most distal promoter upstream of *algZ*.

**Figure 6.**
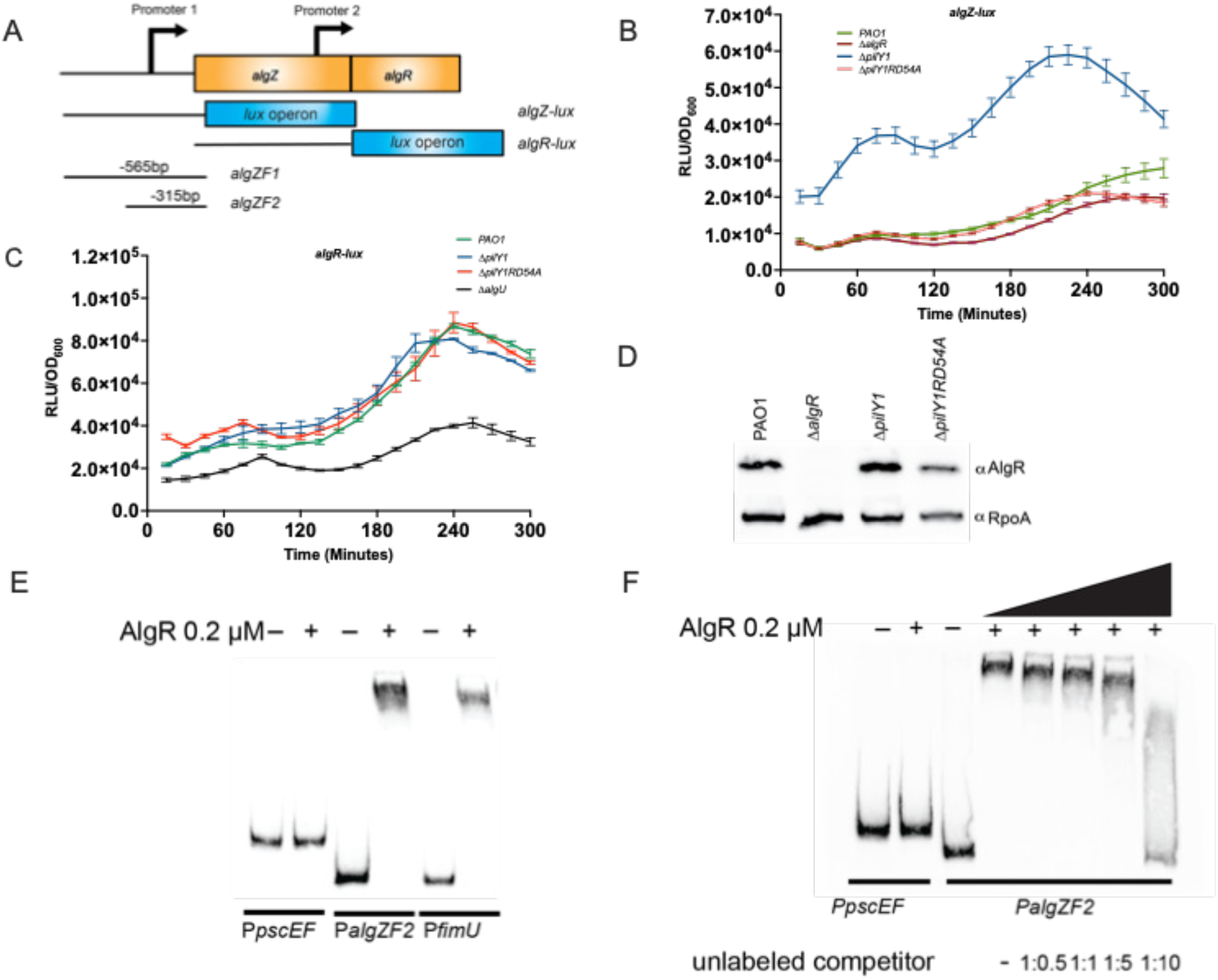
Autoregulation of the *algZ/R* operon. A. Schematic depicting the transcriptional fusions and the EMSA probes used. B. An *algR-lux* transcriptional fusion tested in the pilY1 mutant strains. C. An *algZ-lux* transcriptional fusions tested in *pilY1* mutant strains. D. Western blot confirming increased AlgR in Δ*pilY1* strain and decreased AlgR in the Δ*pilY1*RD54A strain. E. EMSA showing AlgR binding to the *algZ* promoter region. F. Competition EMSA showing specificity of AlgR binding to *algZ* Fragment 1. G. Competition EMSA showing AlgR specificity for *algZ* fragment 2. Transcriptional fusions were performed in triplicate at least three times. Western blotting was performed four times. A representative western is shown. EMSA9s were performed at least three times. A representative EMSA is shown.

Western analysis was used to confirm increased AlgR levels in the *ΔpilY1* mutant (Fig 6D). AlgR was increased in the Δ*pilY1* strain compared to the wild-type strain PAO1 (Fig 6D) indicating that *algR* is increased in *ΔpilY1*. However, when the Δ*pilY*1RD54A strain was tested, AlgR levels were decreased compared to the Δ*pilY1* parental strain (Fig 6D). A Δ*algR* strain was used as a negative control and RpoA was used as a loading control. These results suggested that increased expression of the *algZ/R* operon results in increased AlgR levels.

To determine if AlgR activation of the *algZ* promoter was direct, we performed gel shift analyses using biotinylated probe fragments representing the promoter region upstream of *algZ*. Purified AlgR (0.2μM) was able to shift both a fragment containing 565 bp upstream sequence as well as a shorter fragment consisting of 315 bp of upstream sequence (Fig 6E-G). We were able to inhibit AlgR binding using increasing concentrations of the unlabeled probes for both algZF1 and algZF2 probes. Interestingly, the algZF1 fragment was competed using lower ratios of unlabeled to labeled probe than the algZF2 fragment Fig 6F and G). These results indicated that AlgR specifically binds to the *algZ* promoter.

### PilY1 and AlgZ/R control PA4781 expression

PilY1 signals through the diguanylate cyclase SadC to increase c-di-GMP levels (20) and this could explain why *ΔpilY1* strains have decreased c-di-GMP. However, it is possible that other genes controlled by PilY1 might also be involved in c-di-GMP metabolism. From our RNAseq analysis, we hypothesized that PA4781 might contribute to the decreased c-di-GMP levels in the Δ*pilY1* mutant. PA4781 was previously suggested to degrade c-di-GMP (78), but another study suggested that PA4781 has another role (79).

We first wanted to determine how PilY1 controls PA4781. We constructed a PA4781 transcriptional fusion to further determine the regulation of PA4781 (Fig 7A and B). PA4781 expression was increased in a Δ*pilY1* mutant confirming the RNAseq data. Complementation of the Δ*pilY1* mutant with a single copy of *pilY1* and induction of 0.1% arabinose (or 0.5%) was able to fully restore levels to the wild-type after 120 minutes (Fig 7B). Therefore, PilY1 is important in controlling the expression of PA4781.

**Figure 7.**
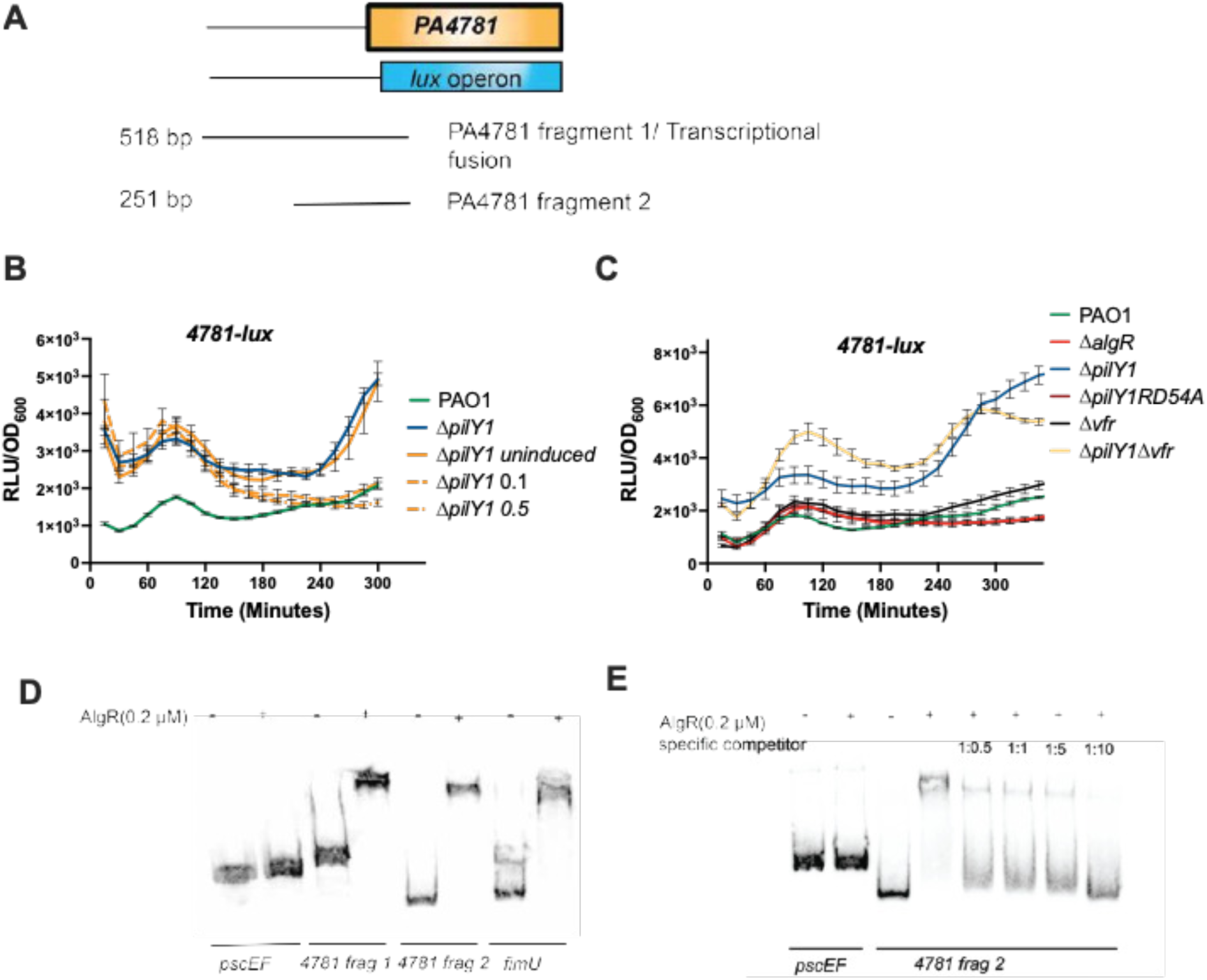
*PA4781* expression. A. Schematic depicting the *PA4781* transcriptional fusion and fragments of the *PA4781* promoter region used for EMSA analysis. B. Complementation of PilY1 reduces *PA4781* expression. A *PA4781* transcriptional fusion was tested in a complemented strain consisting of *pilY1* under an arabinoseinducible promoter. Percentages to the right (0.1 and 0.5) indicate the percentage of arabinose used. C. Testing the *PA4781*-lux fusion to demonstrate AlgRdependence. D. EMSA demonstrating that AlgR binds to the PA4781 promoter region. E. Competitive EMSA showing specificity of AlgR binding to the *PA4781* promoter. Transcriptional fusions were performed in triplicate at least three times. EMSA9s were performed at least three times. A representative EMSA is shown.

We also analyzed the PA4781-*lux* fusion in strains to understand how PA4781 is controlled in the context of PilY1. Because the AlgZ/R system and Vfr are important regulators in the Δ*pilY1* background (19), we investigated these regulators. When the allele of *algR* that encodes a phosphorylation-incompetent form of AlgR in the *pilY1* mutant was tested (Δ*pilY1*RD54A), PA4781 expression was decreased from *ΔpilY1*, suggesting that phosphorylated AlgR is required for activation of the PA4781 promoter (Fig 7C). A Δ*algR* mutant was also tested and had similar expression levels to the wild-type strain. We also tested this fusion in the Δ*vfr* and Δ*pilY1*Δ*vfr* double mutant to determine if Vfr might also regulate PA4781 expression. Mutation of *vfr* did not play a role in regulating PA4781 compared to the wild-type (Fig. 7C). The *ΔpilY1*Δ*vfr* mutant had similar PA4781-*lux* activity as the *ΔpilY1* mutant, suggesting that increased PA4781 expression is not due to Vfr. These data suggest that PilY1 prevents the AlgZ/R system from activating PA4781 expression and that phosphorylated AlgR is required for transcriptional activation.

To determine if AlgR directly regulates PA4781, we performed gel shift analyses. Biotinylated PCR products were used to determine if AlgR directly bound to the PA4781 promoter region. A putative AlgR-binding site was located 425bp upstream of PA4781. Two probes were made and incubated with 0.2μM of purified AlgR (Fig 7A). A probe containing 581 bp of upstream sequence bound purified AlgR that contained a potential AlgR-binding site. A shorter probe encompassing 251 of upstream sequence also bound AlgR, even though no AlgR consensus sequence was found in this region (Fig 7D). Competition experiments with the smaller upstream fragment using unlabeled probe were able to reduce AlgR binding indicating a specific interaction of AlgR with the shorter promoter fragment (Fig 7E). From these data we conclude that AlgR directly controls the expression of PA4781. Altogether, PilY1 decreases PA4781 expression by preventing AlgR activation.

### PilY1 is does not contribute to fitness in an acute pneumonia model

Several studies have indicated that *pilY1* mutants are attenuated (11, 33, 34, 80). However, only one of these studies used a mouse model. The strain used was an uncharacterized clinical isolate and the bacteria were encased in alginate beads to model a chronic infection (33). We tested the Δ*pilY1* strain derived from the PAO1 background to determine the virulence of this strain in an acute pneumonia model using competitive infections. Both PAO1 and the Δ*pilY1* strains were labeled using either a gentamicin resistance cassette (PAO1) or a tetracycline resistance cassette (Δ*pilY1*). The PAO1 strain was mixed in a 1:1 ratio with the Δ*pilY1* strain at a dose of 1 x 10*^7^* CFU/mouse (a lethal dose at 24 hours). We expected the *ΔpilY1* mutant to be less competitive to the wild-type based on previous studies. After 16 hours of infection, the lungs, spleen, and liver were homogenized and plated on differential media to distinguish between the wild-type and mutant and calculate the competitive index (Fig 8). Blood was also analyzed; however, there were no bacteria detected in the blood from 4 mice. Results showed that the Δ*pilY*1 mutant outcompeted the wild-type as indicated by CI values greater than 1.0 in the lung (CI= 1.63) and blood (CI=5.9) (Fig 8). However, in the liver and spleen, the CI was close to 1 indicating that the Δ*pilY*1 mutant was able to compete as well as PAO1 in these tissues (CI= 1.15 and 1.1 respectively) (Fig 8). Overall, these results suggest that the Δ*pilY*1 mutant is more competitive in certain tissues and at least as competitive as the wild-type strain using the acute pneumonia model.

**Figure 8.**
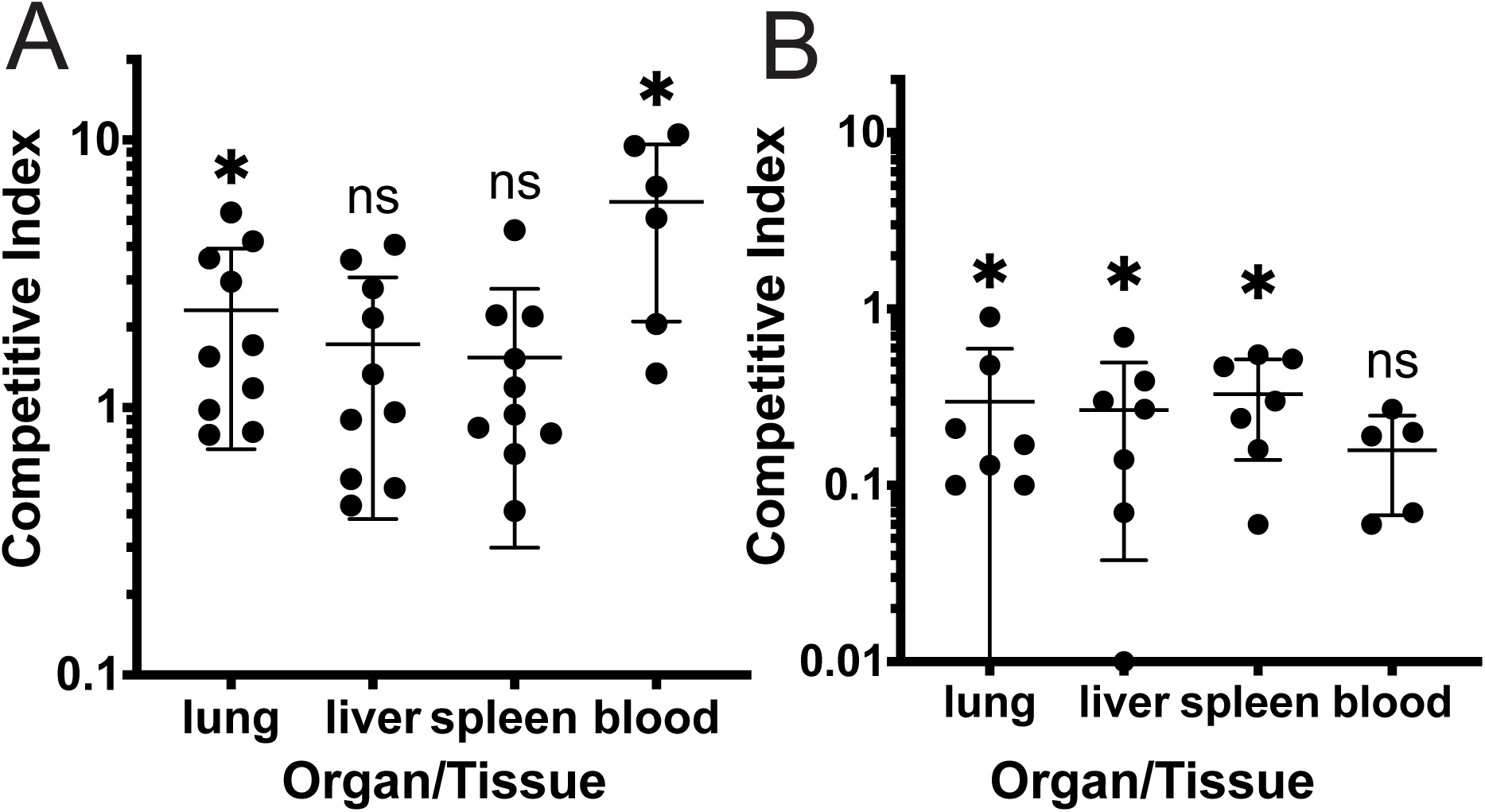
Assessment of PilY1 in *P. aeruginosa* virulence in an acute pneumonia model. A. Competitive index of Δ*pilY1* versus PAO1 during infection. B. Competitive index of Δ*pilA* versus PAO1during infection. Mice were infected for 16 hours via oropharyngeal aspiration. Each point represents a competitive index. n=6-10, One mouse had no detectable bacteria in blood (Δ*pilA*/PAO1). Statistical analysis was using the Wilcoxon signed rank test. *p*< 0.05

The type IV pili mediate close interaction of bacteria with host cells in in vitro studies, but few studies have tested if type IV pili are important in pathogenesis in vivo. We tested a Δ*pilA* mutant in a competition experiment as described for Δ*pilY1*. In contrast to the Δ*pilY1* data, a Δ*pilA* mutant was attenuated in the lung, liver and spleen (Fig 8B). The CI of Δ*pilA* was not significantly different in the blood, but this was likely because we were only able to enumerate CFU’s from 5 mice. Altogether, these data indicate that a Δ*pilA* mutant is attenuated in the acute pneumonia model.

## Discussion

T4P attach many bacterial pathogens to the host (81, 82) and are also important in relaying environmental cues into the bacteria cell (83). *P. aeruginosa* uses the minor pilin, PilY1, as the major adhesive component of the T4P and is important in mechanosensing and virulence (11, 19, 33, 34, 84). Determining what genes PilY1 controls, could shed light on how *P. aeruginosa* interacts and survives within a host. Here, we demonstrate the importance of PilY1 in controlling second messengers and identify new targets PilY1 regulates. We found that PilY1 is important in regulating the major adenylate cyclase, *cyaB*. An unannotated putative cyclic di-GMP phosphodiesterase, PA4781, was also controlled by PilY1. Both *cyaB* and *PA4781* are controlled by the AlgZ/R two-component system illustrating the importance of PilY1 repressing the AlgZ/R system. Additionally, we demonstrate that PilY1 plays a nuanced role in virulence in the acute pneumonia model. PilY1 is necessary for proper cAMP production by preventing AlgZ/R activation of *cyaB*. Overall, our study increases our understanding of how PilY1 and the T4P aid *P. aeruginosa* in sensing the environment.

A major finding of our study revealed that PilY1 controls cAMP levels. It does so by preventing the phosphorylation of AlgR thereby preventing *cyaB* expression. Other pili components have been shown to regulate CyaB activity (12, 23, 53, 54, 85), but none of the pili components have been shown to control the expression of *cyaB*. RNAseq analysis suggested that the slight increase in cAMP was due to the increased expression of the major adenylate cyclase, *cyaB*. We determined that the AlgZ/R system is responsible for increasing *cyaB* expression in the *ΔpilY1* strain. Because PilY1 prevents AlgZ/R activity (19, 34), PilY1 has a major role in regulating cAMP levels. Using transcriptional fusion analyses, gel shift studies, and site-directed mutagenesis, we firmly establish that the AlgZ/R system regulates cAMP under some conditions. This control of cAMP levels is important to allow bacteria to move from a planktonic lifestyle to a biofilm lifestyle (31, 32). Other pili components control the activity of CyaB after expression (12, 19, 54); but this would occur after PilY1 has bound to a surface. It may be that there is expression of *cyaB* in the planktonic phase, but the CyaB enzymatic function is not active until the T4P extend and contract after binding a surface. This would help explain the surface activated virulence seen previously (11).

The role of PilY1 in virulence is still unclear. Previous studies have indicated that PilY1 is important in virulence (11, 33, 34). Only one of the previous studies used a mouse model of infection. This study used a chronic isolate and chronic infection model using alginate beads and only examined the lungs. However, the authors also found an increase in CFU’s of the *ΔpilY1* strain compared to the wild-type in a competition experiment (33). The competitive index (CI) is a sensitive measure of comparing virulence between mutant and wild-type strains (86). We found that the *ΔpilY1* strain was at least as fit in the liver and spleen and more fit than the wild-type in the lungs and blood. While we did not evaluate the immunopathology during these infections, the fact that the host is incapable of eliminating the *ΔpilY1* strain complicates assigning the *ΔpilY1* strain as attenuated. It may be that the timepoint used in the acute pneumonia model of 16 hours was not sufficient to see a difference between the two strains. For instance, if the *ΔpilY1* strain caused more neutrophil infiltration, this could result in more pathology early on, but later timepoints might find that the mutant strain is eliminated at greater numbers than the wild-type.

A previous study indicated that a *pilA* mutant was attenuated in a survival study (9). Our work confirms this result using a slightly different model and a different strain. Using oropharyngeal aspiration instead of intranasal inoculation to introduce bacteria ensures more bacteria in the lungs (87, 88). This makes our model relevant to the study of ventilator-associated pneumonia. A recent transcriptomic analysis of a Δ*pilA* strain suggested that the main reason for attenuation is due to cAMP production (89). It is possible that the increased cAMP in the Δ*pilY1* is responsible for the increased virulence. Given the importance of careful regulation of second messengers, inappropriate levels at a given time may attenuate *P. aeruginosa* virulence. Additionally, our study has demonstrated that while some pili components are required for virulence, such as PilA, others such as PilY1, can prevent growth in certain tissues.

Based on our data and the work of others, we propose that the phosphorylation status of AlgR is what determines whether Vfr and AlgR act synergistically or in an opposing fashion (Fig. 9). In the case of *pilY1* expression, phosphorylated AlgR and Vfr activate the *fimU* operon (which includes *pilY1*) (21, 22). The work presented here suggests that the AlgZ/R system can also activate *cyaB* expression allowing cAMP production that can then activate Vfr.

**Figure 9.**
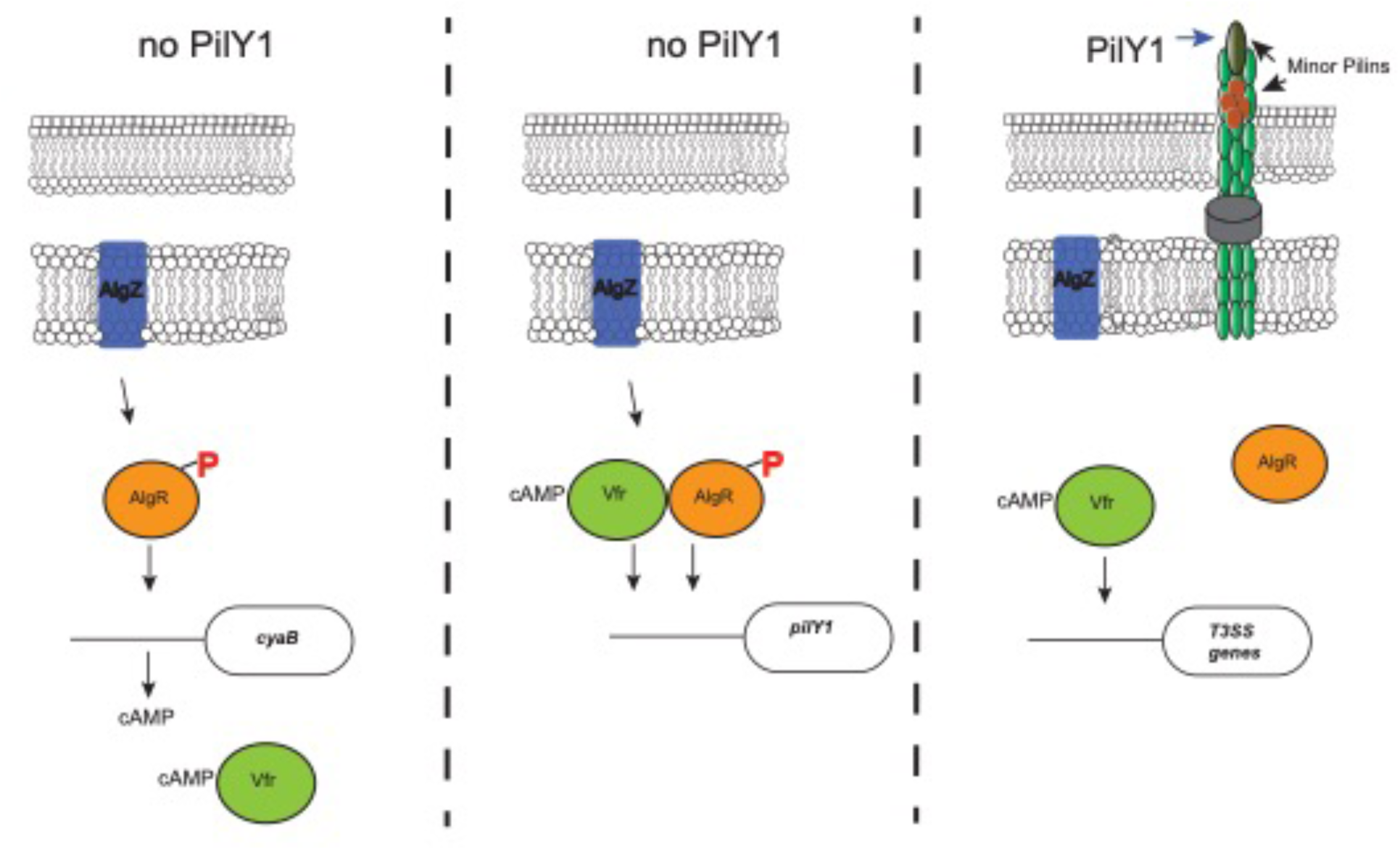
PilY1 exerts its effects by turning off the AlgZ/R system and is required for proper virulence gene coordination. Model depicting how PilY1 controls AlgZ/R and regulates AlgR and Vfr activity. Left side shows that AlgZ/R activates expression of *cyaB* so that cAMP can be made and activate Vfr. Middle panel shows that phosphorylated AlgR and activated Vfr activate expression of the *fimU* operon encoding *pilY1*. Right side shows that PilY1 turns <off= the AlgZ/R system and activated Vfr can activate the T3SS genes.

Phosphorylated AlgR and Vfr can then activate the *fimU* operon (including *pilY1*). After expression of the *fimU* operon, PilY1 turns off the AlgZ/R system. This would then allow Vfr to activate other targets such as the T3SS. Given that AlgR can inhibit T3SS expression (90) and that the T3SS is activated via surface contact (91, 92), PilY1 deactivation of AlgZ/R is required for Vfr to activate the T3SS. Could phosphorylated AlgR be used to prevent Vfr from activating the T3SS expression? This could explain the reduced virulence of *algR* overexpressing strains previously described (93). In addition, does PilY1 continue to turn off the AlgZ/R system when PilY1 is engaged with its cognate ligand on a host cell? Further studies using *pilY1* variants instead of deletion mutants are required to answer these questions.

PilY1 has garnered attention for its role in mechanosensing and virulence. In this report, we have identified an important role of PilY1 in controlling cAMP as well as c-di-GMP. This aids our understanding of how the T4P can engage in signaling but also suggests that PilY1 function is important in other locations as well. We also have aided in the understanding of how different regulatory pathways intersect. Vfr and AlgZ/R can work together to activate expression of the *fimU* operon. Once *pilY1* is expressed, PilY1 turns off the AlgZ/R system and other pili components can activate CyaB allowing Vfr to activate expression of virulence genes. Further work is necessary to unravel the role of PilY1 in *P. aeruginosa* pathogenesis. Understanding signal transduction networks can identify new therapeutic targets and allow new ways to treat *P. aeruginosa* and other antibiotic-resistant bacteria.

